# What are the most influencing factors in reconstructing a reliable transcriptome assembly?

**DOI:** 10.1101/220269

**Authors:** Noushin Ghaffari, Jordi Abante, Raminder Singh, Philip D. Blood, Lenore Pipes, Christopher Mason, Charles D. Johnson

## Abstract

Reconstructing the genome and transcriptome for a new or extant species are essential steps in expanding our understanding of the organism’s active RNA landscape and gene regulatory dynamics, as well as for developing therapeutic targets to fight disease. The advancement of sequencing technologies has paved the way to generate high-quality draft transcriptomes. With many possible approaches available to accomplish this task, there is a need for a closer investigation of the factors that influence the quality of the results. We carried out an extensive survey of variety of elements that are important in transcriptome assembly. We utilized the human RNA-Seq data from the Sequencing Quality Control Consortium (SEQC) as a well-characterized and comprehensive resource with an available, well-studied human reference genome. Our results indicate that the quality of the library construction significantly impacts the quality of the assembly. Higher coverage of the genome is not as important as the quality of the input RNA-Seq data. Thus, once a certain coverage is attained, the quality of the assembly is mainly dependent on the base-calling accuracy of the input sequencing reads; and it is important to avoid saturating the assembler with extra coverage.

## I. INTRODUCTION

Every living metazoan is a collection of cells with different functionalities. In order to have a thorough understanding of molecular level interactions of a given species, knowledge of the arrangement of its genome (DNA) and transcriptome (RNA) is key. In the past decades, efforts to uncover the underlying genetic and transcribed material of organisms has spanned many species, including Homo sapiens, Mus musculus, Drosophila melanogaster, Danio rerio, and Arabidopsis thaliana, which have expanded our awareness and ability to develop clinical, environmental and economical gene targets. Ascertaining genome of non-model organisms has been a rapidly expanding research direction in the light of improved sequencing technologies.

Sequencing pertains to determining the sequence of DNA/RNA fragments using different methods. Ultimately, leading to gene recognition and making consensus genomes for non-model species, which need further steps to become the reference genome. The modern sequencing technologies pioneered by Sanger sequencing methodology in 1970s [1] and continued to move forward with the introduction of Next Generation Sequencing (NGS) techniques [2]. Yet, regardless of the technology, sequencing methods shear (intentionally or through manipulation) the input DNA/RNA material, which causes disjointed pieces representing the original genome, with the caveat of being much shorter than source material. The resulting pieces must be connected to each other to retrieve the original genetic material.

Accodingly, there can be an immense computational demand in generating a such a reference genome or transcriptome. There has been significant improvements in the algorithms that take the sequencing outputs and generate a de novo consensus genome. This process is referred to as genome/transcriptome assembly [3]. In the last decade, the next generation sequencing technologies dominated the sequencing field, thus, most of the efforts in the bioinformatics community concentrated on NGS-based tools, e.g. Trinity [3], SoapDenovo [4], ALLPATHS-LG [5]. Although assembling the references is important, ensuring the accuracy of the consensus is even more critical. Reporting genomes with avoidable shortcomings can create many future issues. This current study aims at reconstructing human transcriptome and investigating the factors that affect the quality and accuracy of the reported consensus. To that end, high quality sequenced RNA, well-annotated reference genome and highly reliable assemblies are essential. The US Food and Drug Administration (FDA) has coordinated the Sequencing Quality Control project (SEQC/MAQC-III) with the goal of assessing the technical performance of RNA-Seq experiments comprehensively. The SEQC consortium generated benchmark datasets of well-studied reference samples, sequenced at multiple sites, and using different sequencing platforms, with controlled settings. The generated RNA-Seq data is used in separate studies to measure quality metrics, spike-in controls, limits of detection, effects of analytic pipeline and assessments of RNA-Seq accuracy and reproducibility. Samples were distributed among six independent centers in the study: 1- Australian Genome Research Facility (ILM1), 2- Beijing Genomics Institute (ILM2), 3-Weill Cornell Medical College (ILM3), 4- City of Hope (ILM4), 5- Mayo Clinic (ILM5) and 6- Novartis (ILM6). The ILM2, ILM3 and ILM5 were selected as the official sites. The SEQC used the experimental design and commercially available samples from Microarray Quality Control Consortium MAQC I [6], [7]. Sample A is the well-characterized Universal Human Reference RNA (UHRR), and sample B is Human Brain Reference RNA (HBRR). The synthetic RNA from the External RNA Control Consortium (ERCC) [8] was spiked in. Samples C and D were generated by mixing samples A and B in ratios of 3:1 and 1:3, respectively. Each one of samples A and B had 5 replicates. Replicates 1 to 4 were prepared in each site. The vendor prepared the fifth replicate. To examine the effect of the instrument on the RNA-Seq experiments, all the samples were sequenced using Illumina HiSeq 2000, and for generating longer reads three sites sequenced samples A and B using the Roche 454 GS FLX platform. The other next generation sequencing technology, SOLiD, was also used. The SEQC consortium overall sequenced 108 libraries on a HiSeq 2000, 68 libraries on SOLiD, and 6 libraries on a Roche 454, generating more than a 100 billion reads, for samples A to D.

The first study that focused on RNA-Seq assessments was recently published [9], systematically examining the impact of site-specific bias in detecting differentially expressed genes, using different RNA-Seq analysis methods. They showed that none of the tested technologies provided reliable absolute quantification and relative expression measures that agreed well across validation platforms: RNA-seq, qPCR and microarrays. However, the built-in design of the study was satisfied by the majority of genes. They observed sensitivity of results to analysis pipeline choice, and provided a suggestion for sequencing depth: An effective sequencing depth is clearly contingent on the experimental goals, with simple gene-level expression profiling only requiring 550 million single-ended reads for an appropriate analysis pipeline [9]. The RNA-Seq mapping and differential expression testing pipeline such as TopHat2 and CuffDiff [10], Magic [11], BitSeq [12], Subread [13] and r-make incorporating STAR [14] were examined and their performance was compared. In addition, the SEQC reference datasets were an invaluable resource for a systematic characterization of measurements, and for making reliable conclusions from large-scale experiments.

The availability of SEQC datasets provides an excellent opportunity for the scientific community to examine the RNA-Seq experiments, and to learn more about its variety of characterizations and applications. The SEQC consortium has focused on RNA-Seq analysis pipelines for differential expression detection, splice junction discovery, sample differences, etc. However, the performance of different SEQC RNA-Seq datasets has not been evaluated for transcriptome assembly (to the best of our knowledge). It is crucial to understand the performance of transcriptome assemblies to improve current practices. Understanding the factors that affect transcriptome assembly is also very important. To achieve these goals, reliable input data, which were generated with the highest standards, are necessary. Here, we examine the SEQC RNA-Seq data for assembling human transcriptome. The comparison of our resulting de-novo assembled transcriptomes with the well-annotated human transcriptome provides insights to the pros and cons of the assembly procedure. The current study aims to generate assemblies, with fixed parameters, to examine the effects of built-in settings of SEQC data on the results.

## II. METHODS

This study examined the effects of different sequencing sites, samples and sequencing depths on the assembled transcriptome, using well-characterized RNA-Seq data from SEQC. The study encompasses the following three main aspects:

- Aspect I: Site effect examinations
- Aspect II: Coverage/Sequencing depth effect examinations
- Aspect III: Library effect examinations

Aspect I pooled four site-prepared replicates by six Illumina centers (samples A and B separately) and proceeded to the transcriptome assembly. Since the fifth replicate was prepared by the vendor, it was excluded from this aspect. Including only the replicate prepared by each site enabled us to examine the effect of library preparation and sequencing in each individual center. The other objective of this aspect was studying the impact of the coverage. Aspect II evaluated the effect of sequencing depth on the transcriptome quality. All the replicates for sample A (and B) from all the sites were pooled, and then assemblies were constructed using 25%, 50%, 75% and 100% of the data. The assemblies of Aspect II demanded tremendous computational power due to their very high genome coverage. In Aspect III, three official Illumina sites participated. The goal was to compare the vendor-prepared and site-prepared samples. For each site, we compared a random sample of the pooled library with the vendor-prepared library assigned to that center by sampling the same number of reads as in the latter to achieve comparable coverages. In addition to three main aspects mentioned above, we re-examined Aspect I and Aspect II with selected input data. The objective of these aspects was to determine the influence of in-silico normalization option on the assembly quality. We will refer to these two assemblies as Aspect I-Norm and Aspect II-Norm.

In order to examine all three main objectives of the study (and additional two aspects), we designed a multi-step workflow, depicted in Figure 1. There are three major components: 1- pre-processing, 2- assembling the transcriptome, 3- post-processing.

**Fig. 1:**
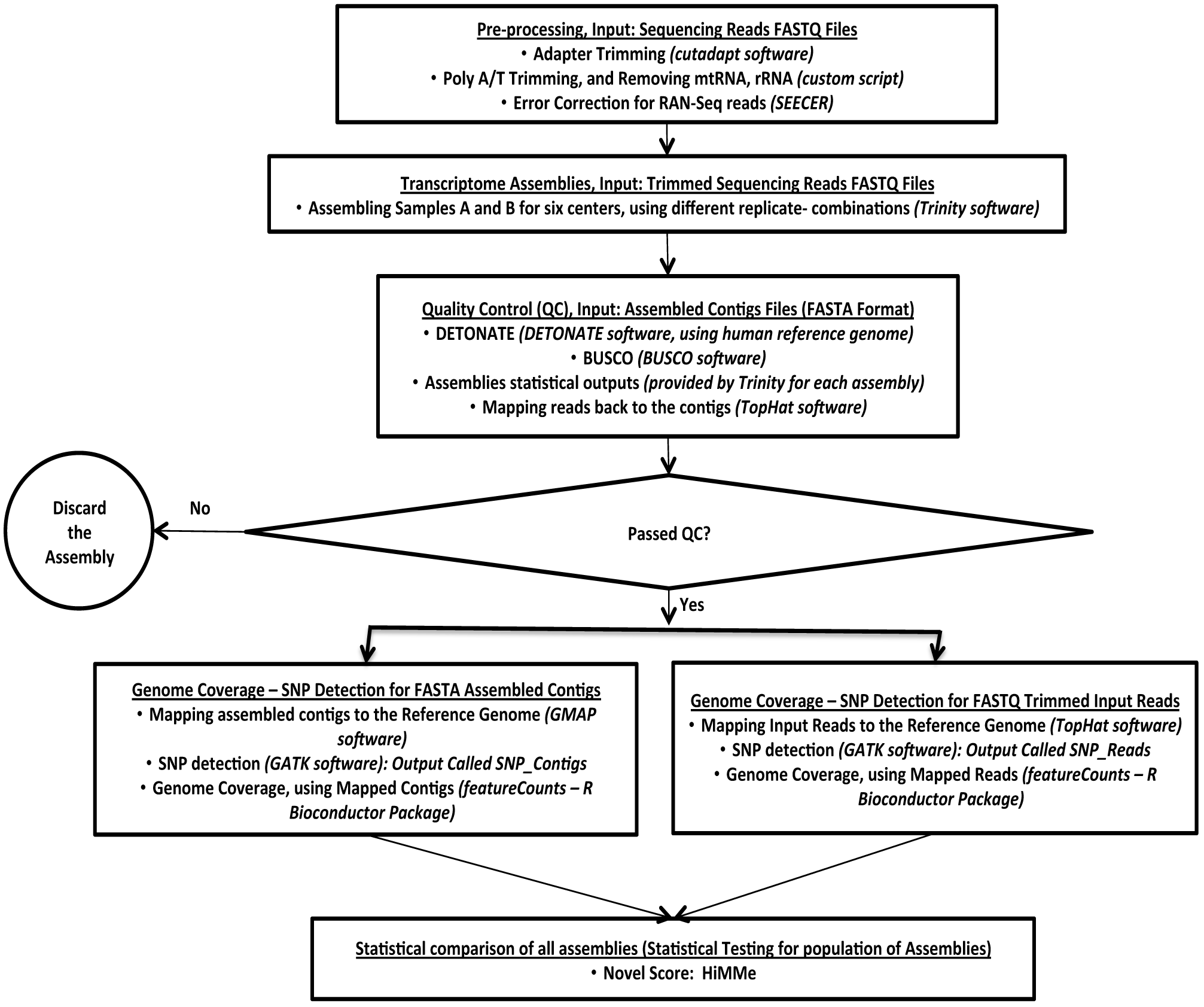
Study plan. This workflow flowchart shows the steps and tools that are used for the current study.

The pre-processing phase started by discarding any reads with attached adapter sequences. For adapter trimming we used Cutadapt [15]. In order to remove poly A/T tails we used Flexbar [16]. The RNA-Seq tends to include sequenced fragments (reads) with ribosomal RNA (rRNA) and the mitochondrial chromosome. We eliminated the rRNA and mitochondrial reads that mapped to their respective references. As for sequencing errors, SEECER [17] is a powerful tool that corrects errors RNA-seq data. In the last step of our pre-processing we employed SEECER on the remaining reads.

The most computationally intensive part of our workflow was the transcriptome assembly. All the fragments that successfully passed the pre-processing stage were used to construct the respective transcriptome assembly. To that end, we used Trinity [3], a state-of-the-art *de novo* assembly algorithm developed specifically for reconstructing transcriptomes based on *de Bruijn* graphs. Transcriptome assembly is challenging mainly because RNA-Seq data coverage levels are not evenly distributed. Furthermore, alternative splicing complicates assembly from individual genes. The goal of Trinity is to deliver one graph per expressed gene. This assembly tool consists of three parts: 1) Inchworm, 2) Chrysalis, and 3) Butterfly. During these three steps, Trinity makes linear contigs from RNA-Seq reads, generates and expands de Bruijn graphs, and finally outputs the transcripts and isoforms. There are many transcriptome assembly tool available, however, the intention of this study was not to compare assembly algorithms, but rather to generate transcriptomes with a reliable pipeline that would highlight the effects of Aspects I-III.

The SEQC data was generated at multiple sites and included well-studied samples. Samples A and B differ since sample A (UHRR) consists of ten pooled cancer cell lines, and sample B (HBRR) is from multiple brain regions of 23 donors. Our objective was to examine the effects of different sites, samples and sequencing depths on the assembled transcriptome. We utilized samples sequenced by the Illumina platform. We selected samples A and B (sample-type effect), across all six sequencing site (site effect), and assemblies were done for different number of pooled replicates (sequencing depth effect). There are four replicates available for samples A and B, sequenced in each center. The fifth replicate is prepared by the vendor and sequenced in each site. The examination of all transcriptome assemblies provided insights on successful RNA-Seq experiments and transcriptome assemblies.

In the post-processing stage, each assembly went through a comprehensive pipeline of quality measurement tools. The statistics of assemblies were used to assign a ranking to the result, e.g. between similar assemblies, the one with larger N50 will receive higher ranking. Additionally, mapping results using GMAP [18], and SNP detection process using GATK [19] are important for ranking the assemblies. The common assembly statistics, such as maximum contig length and N50 were used to determine the quality of the results, and to compare the assemblies using different parameters. However, as other studies noted [20], [21], in our recent paper [22] we showed that using additional measures for quality control and comparison of assemblies is important. Relying only on the assembly statistics can be misleading and result in overstating the power of the assemblers. Therefore, we included the following steps in our validation/assembly comparison pipeline: examining the statistics of each assembly and ranking assemblies accordingly, comparing completeness of the assemblies with Benchmarking Universal Single-Copy Orthologs (BUSCO) [23], and assigning scores to the assemblies using DETONATE (DE novo TranscriptOme rNa-seq Assembly with or without the Truth Evaluation) [24]. The latter uses both the reference genome as well as the input RNA-seq reads, to compute multiple metrics and probabilities to assess how well supported the assembly is in light of the previous knowledge. Employing all these quality control (QC) metrics and comparison/ranking tools ensures the quality of the outputs, and will assist us to find the differences between assemblies based on samples A and B sequenced in different sites with different coverage. Furthermore, we applied our recently developed algorithm HiMMe [25] that uses the genetic content of assemblies as a proxy for assembly reliability. This novel method is based on hidden Markov model (HMM) theory, and computes an individual score for each contig by looking at the k-mer transitions observed, as well as a global score for the assembly. The HiMMe algorithm is trained using previous knowledge about the samples. In this study, the reference transcriptome and a SNP database were used to learn the model [25].

## III. RESULTS

This section presents the most relevant findings related to our three aspects, and the remaining material can be accessed in the supplemental document.

The transcriptome assemblies generated by Trinity were examined with multiple quality control measures. The assembly statistics, provided by assembly software, is an important starting control. The N50 defines the contiguity of the resulting contigs by having at least half of the contigs longer than its value. Figure 2 depicts the N50 values for all objectives of this study for sample A. The corresponding plot for sample B shows similar trends (supplemental document).

**Fig. 2:**
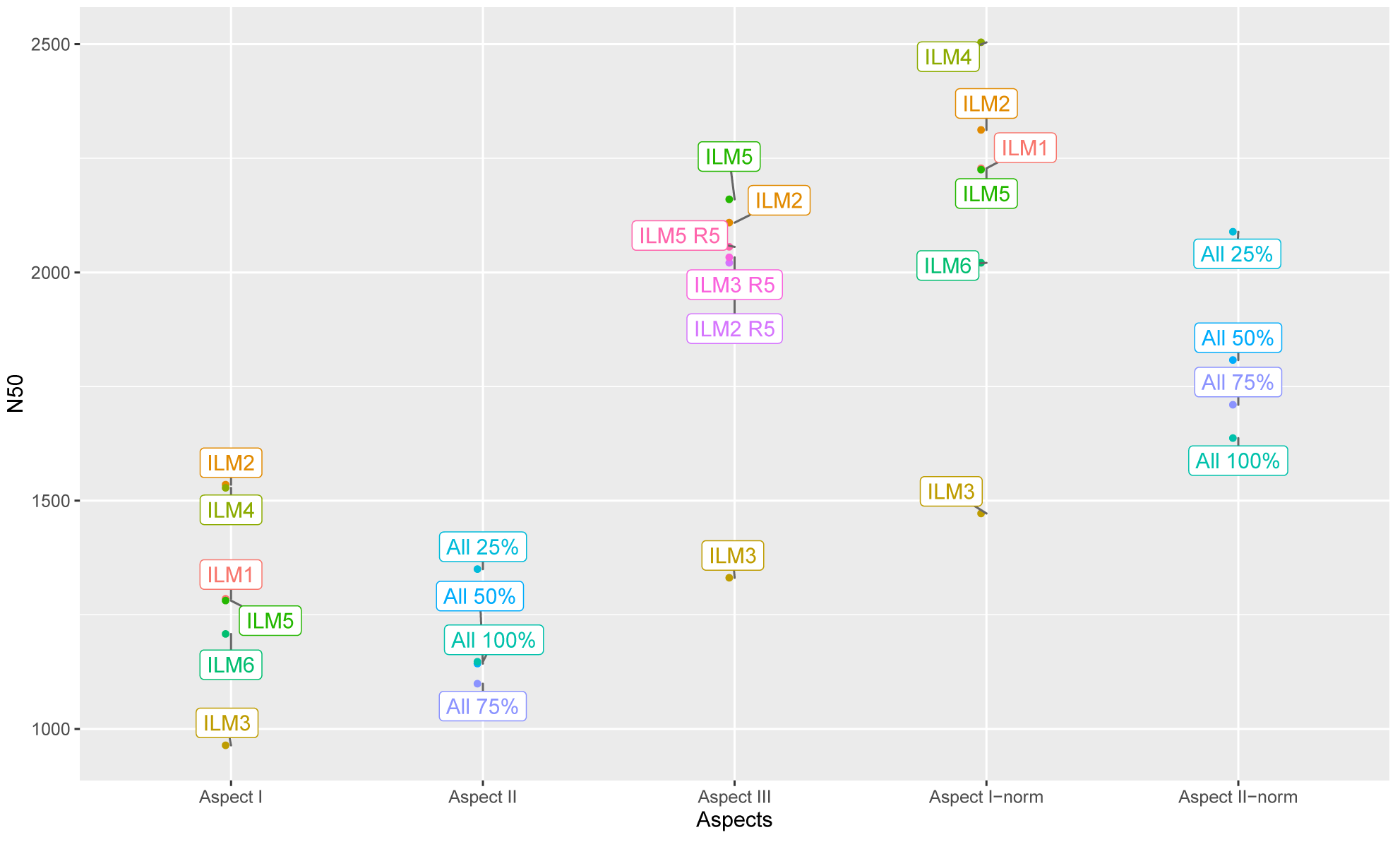
N50 values for the five studied aspects.

We also considered the effect of the input read volume for each assembly. The number of input reads was lower than the original in SEQC data since during the pre-processing step reads with adapter or ploy A/T tails, as well as reads mapping mitochondrial DNA, were eliminated. The genome coverage is computed by dividing the input reads number with the human genome size. The transcriptome size is more variable than genome size due to expression of genes in different times, difficulty in narrowing down coding regions, tissue specification, etc. Thus, this paper uses genome coverage, unless otherwise stated.

Aspect I aims at comparing Illumina sequencing sites with respect to library preparation and the input volume generated, by pooling and then assembling four replicates generated in each site. As Figure 2 shows all sites generated N50 values higher than 1200, except ILM3. Looking at the FastQC [26] plots for different sites, it is apparent that the quality of the libraries in ILM3 site were shifted in GC-content and insert size relative to the rest of the sites. The FastQC plots for SEQC data is provided in the supplemental document. Other studies also noted similar shifts in data QC metrics, with higher per-base error rates generated in ILM3 [27]. Thus, less contiguity measure should be expected and it is visible in our results. The genome coverage ranged from 5x to 35x among 6 sites, including all replicates. Looking at the individual libraries for each replicates, which was sequenced in 8 lanes of a flowcell, the libraries generated varying reads from 1M up to 10M. The high genome coverage for each replicate is achieved by numerous libraries for each replicate. ILM2 and ILM4 had 17x and 19x coverage and generated highest N50 values. ILM6 had the highest 35x coverage and its N50 is moderate, whereas ILM3 with minimum N50 had 9x coverage. These results stress the importance of having quality sequencing reads for producing high quality assemblies.

In Aspect II, we pooled all the replicates for sample A and then randomly picked 25%, 50%, and 75% of the data for assemblies. We utilized the entire pooled data set in the last assembly. The sequencing depth ranged from 40x to 160x from 25% to 100% assemblies with approximately 40x increase in each step. These Trinity assemblies were among the largest ever done at Pittsburgh Supercomputing Center (PSC). Looking at the resulting N50 values for this aspect, increasing the volume of input data negatively influenced the contiguity of the results. The N50 value was highest for 25% run and declined as coverage increased, indicating that overwhelming assembler with too much data will not necessarily improve the quality of contigs. It should be noted that aspect II provides a significant suggestion for assembly with claiming that moderate coverage and high quality data suffices in generating reliable transcriptome assemblies.

Aspect III utilizes replicate 5 prepared by vendor and sequenced in three official sites. As Figure 2 shows, all assemblies with this higher quality library performed well. The sequencing depth for this replicate is 2.7x to 4.6X and still generated the higher N50 values compared with Aspects I and II. In another part of Aspect III, we pooled replicates 1 through 4 prepared at each sequencing center and randomly selected the number of reads to be equal to that of replicate 5. Interestingly, site ILM3, the one with lower quality data, performed poorly, clearly emphasizing on the library preparation effect on assembly quality.

In addition to the three main aspects of the study, we re-evaluated the performance of Aspects I and II using the In silico normalization of RNA-Seq fragments of Trinity [3]. Trinity normalization targets a specified coverage and discards reads that are less useful in assembly. Most transcripts can be saturated with lower coverage and adding more input data only increases the computational cost and may adversely influence the results. The normalization significantly increased the N50 values for Aspects I and II, emphasizing more on the importance of high-quality data and appropriate sequence depth.

The quality of the assemblies was further assessed by the BUSCO tool. As table I shows in aspect I the six Illumina centers performed equally well in recovering BUSCO orthologs. However, ILM3 center had the minimum C measure. In Aspect II the performance is very similar, implying that all runs preserved the appropriate gene information, regardless of their contiguity measurements. The first part of Aspect III, where R5 is used for assembly shows high completeness for the three official site, but the coverage of the second part of the aspect reveals that ILM3 has the lowest BUSCO measurements. The same pattern is observed in other QC steps emphasizing on the less satisfactory results when libraries prepared by ILM3 are used.

**Table I.** The output of BUSCO quality control tool for three primary aspects of the study.

| Aspect | Center/Coverage | Complete (C) | Duplicated (D) | Fragmented (F) | Missing (M) |
| --- | --- | --- | --- | --- | --- |
| Aspect I | ILM1 | 87% | 42% | 4.80% | 7.60% |
|  | ILM2 | 86% | 43% | 5.20% | 8.70% |
|  | ILM3 | 82% | 37% | 8.00% | 9.50% |
|  | ILM4 | 86% | 41% | 4.50% | 8.40% |
|  | ILM5 | 86% | 42% | 5.20% | 7.70% |
|  | ILM6 | 86% | 41% | 5.90% | 7.80% |
| Aspect II | 25% | 87% | 41% | 5.30% | 7.50% |
|  | 50% | 88% | 43% | 4.40% | 7.20% |
|  | 75% | 88% | 43% | 4.30% | 6.90% |
|  | 100% | 88% | 44% | 4.80% | 6.20% |
| Aspect III - R5 | ILM2 | 84% | 42% | 4.70% | 10.00% |
|  | ILM3 | 82% | 39% | 5.50% | 11.00% |
|  | ILM5 | 81% | 39% | 6.10% | 12.00% |
| Aspect III - R1 to R4 | ILM2 | 84% | 43% | 4.80% | 10.00% |
|  | ILM3 | 76% | 35% | 10.00% | 13.00% |
|  | ILM5 | 82% | 39% | 5.40% | 12.00% |

**Fig. 2:**
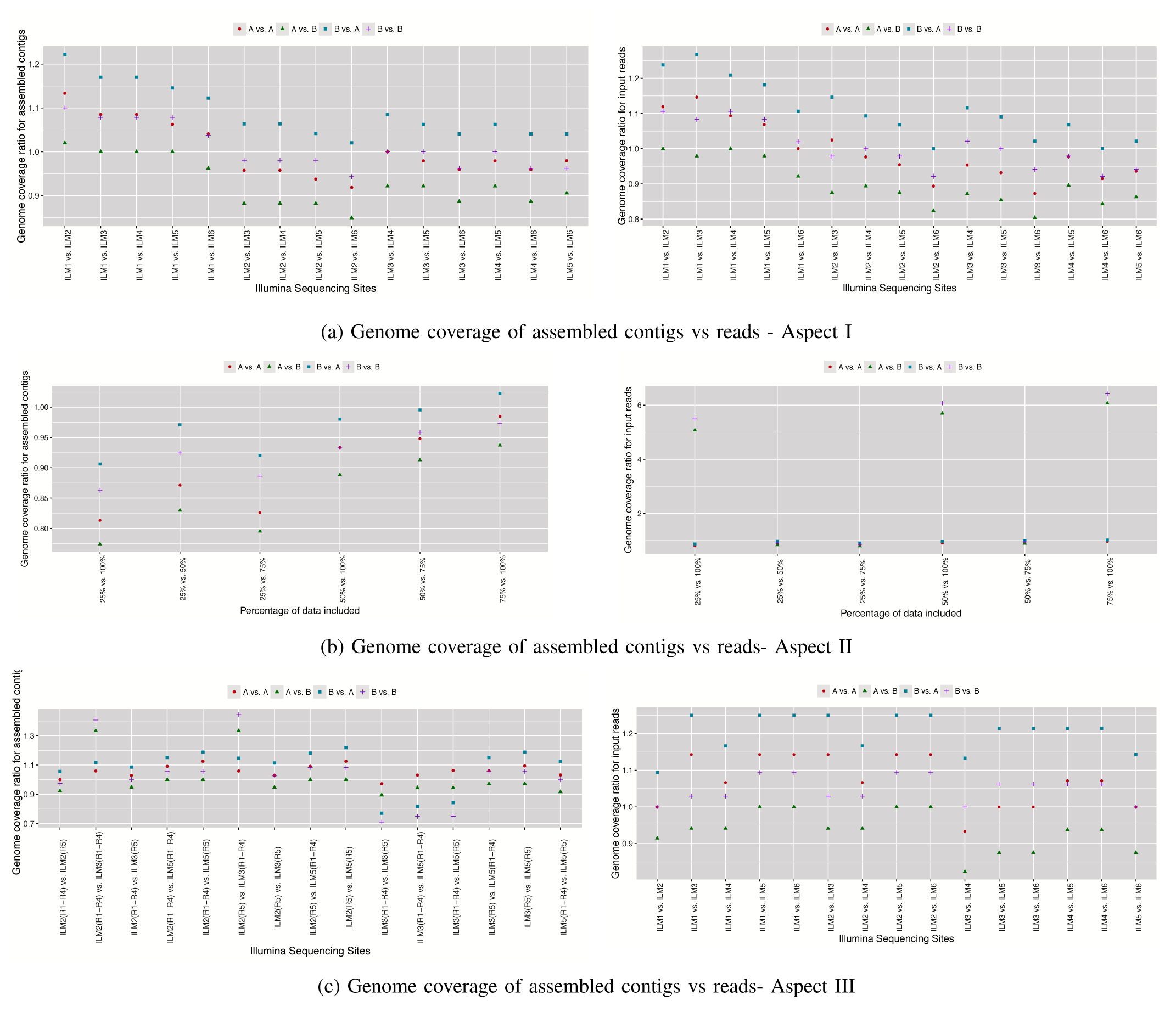
Comparing genome coverage of samples A and B in 6 Illumina sites in Aspects I to III. The value A vs A shows the coverage for sample A in ILMX versus ILMY site. Values A vs B, B vs A and B vs B correspond to the ratios of overages for samples A and B among sites.

Furthermore, we used DETONATE [24] to study the k-mer proportions present in the transcriptome compared to that in the reference. As Table II shows the sites with lower library quality have worse scores, confirming the library preparation quality observations with previous QC measures.

**Table II.** The output of DETONATE for three primary aspects of the study (Sample A).

| Aspect | Center/Coverage | Number of contigs | DETONATE score |
| --- | --- | --- | --- |
| Aspect I | ILM1 | 542,091 | -30,562,340,585 |
|  | ILM2 | 375,745 | -18,347,313,246 |
|  | ILM3 | 604,390 | -22,478,407,936 |
|  | ILM4 | 403,209 | -20,043,442,894 |
|  | ILM5 | 483,867 | -25,860,133,373 |
|  | ILM6 | 535,758 | -35,688,527,059 |
| Aspect II | 25% | 542,321 | -40,252,604,757 |
|  | 50% | 890,995 | -79,325,290,744 |
|  | 75% | 1050484 | -118,717,758,792 |
|  | 100% | 1,091,728 | -157,161,169,416 |
| Aspect III - R5 | ILM2 | 215,811 | -4,838,389,288.10 |
|  | ILM3 | 185,289 | -4,416,597,242.54 |
|  | ILM5 | 161,700 | -3,244,238,513.70 |
| Aspect III - R1 to R4 | ILM2 | 196,240 | -5,052,617,512.70 |
|  | ILM3 | 220,790 | -4,861,137,388.42 |
|  | ILM5 | 157,248 | -3,208,148,228.65 |

The transcriptome reconstruction for the non-model organisms is based on de novo procedures and solely relies on the input sequenced reads. Thus it is difficult to compare the coverage of the resulting transcripts with a reference genome and compare it with the corresponding read coverage. The SEQC homo sapiens samples used here have well annotated reference genome and, in turn, this enabled us with well-annotated reference genome, which enabled us to compare the genome coverage for reads and assembled contigs. Figure 2 shows the relative genome and read coverage for this study. In order to compare the genome coverage, we provided the relative coverage of sample A and B among sites. As the figure shows the contigs and input reads provided similar genome coverage patterns. The consistent trends show the higher coverage of ILM1 among centers and higher relative values for sample B versus sample A. We also observed a lower contig coverage for ILM3 when using replicates 1 through 4.

Additionally, we employed our novel HiMMe method to evaluate all transcriptome assemblies generated. When looking at the score distributions for Aspect I (see Figure 3a), we do not observe major differences between sites in general. However, we do see a significant lower median score for site ILM3 for contigs of length between 10,000-100,000 nucleotides. Thus, we conjecture that the quality of the libraries might have a bigger impact when assembling longer transcripts. When looking at the score distributions for Aspect II (see Figure 3b), we do not observe any trend suggesting that more data improves the output as long as a critical coverage is reached, consistent with previous results. As for the assemblies that belong to Aspect III (see Figure 3c), we do observe a general trend suggesting that the quality of the input data does have an impact on the assembled transcriptome. Transcriptomes assembled using vendor prepared libraries (R5) have a slightly higher median score compared to that of assemblies constructed with pooled reads from R1 through R4, which in turn were prepared by the sequencing sites. As suggested above, this difference becomes larger with longer transcripts. The largest difference is observed in the last bin, where ILM3-R5 performs significantly better than ILM3. These results further the point that the quality of the input reads, once a minimum coverage is secured, is the most determinant factor for reliable transcriptome assemblies.

**Fig. 3:**
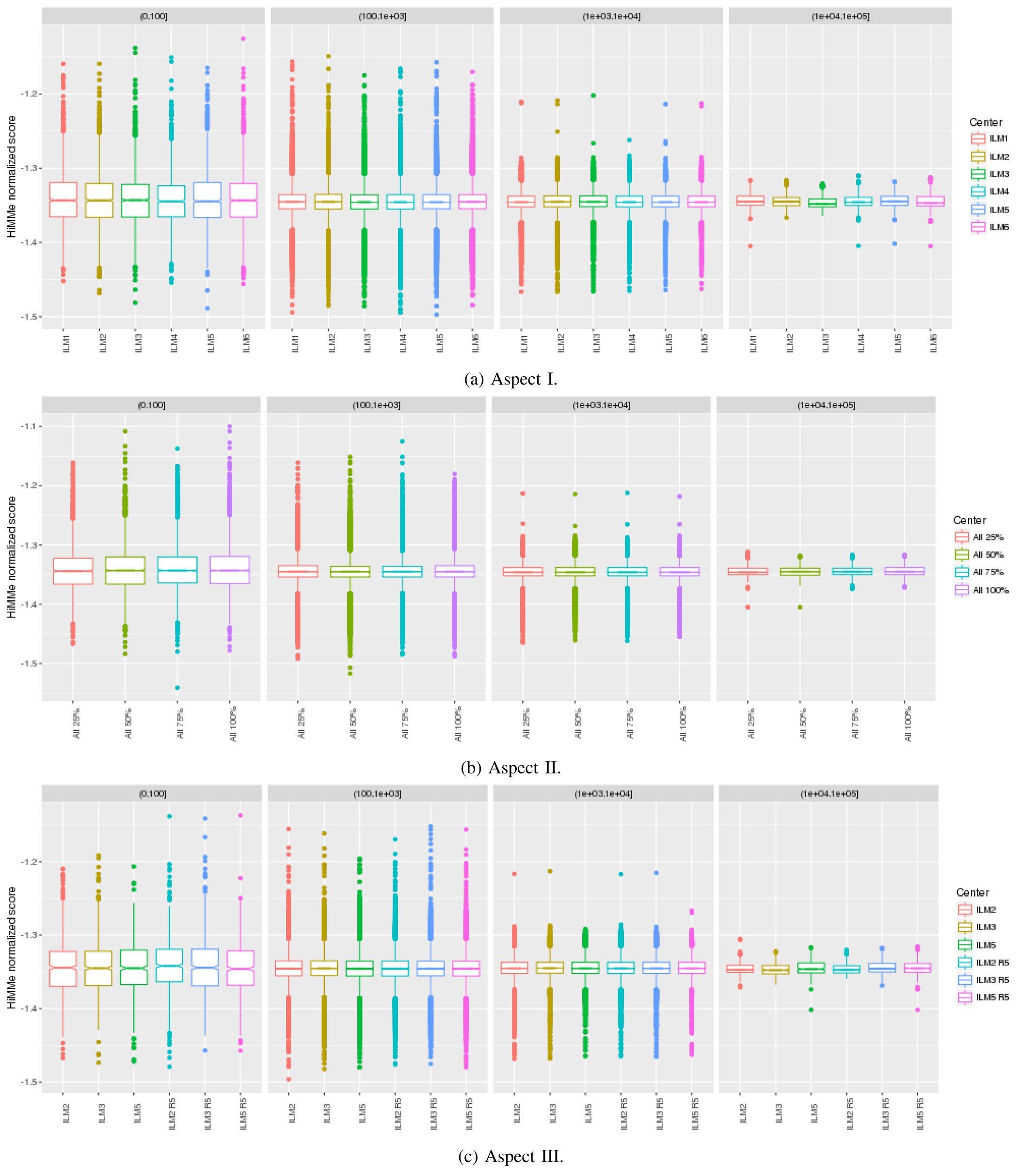
Score distributions for Aspect I, Aspect II, and Aspect III, computed by HiMMe. The scores have been binned, by looking at contig length, in four bins of length ranging between 0-100 nucleotides; 101-1,000 nucleotides; 1,001-10,000 nucleotides; and 10,001-100,000 nucleotides.

For future assessment, we compared the number of called Single Copy Polymorphisms (SNP) that are called by input reads and the assembly results. One should expect considerable differences between the SNPs called by the two sources. Reads are shorter and may have duplication and their alignment to the reference genome capture many more SNPs. In all the three aspects of the study contigs have very close SNP call patterns, and raw reads are clustered nicely. The figure showing these findings can be accessed at supplemental document.

Furthermore, we assessed the quality of each transcrip-tome by calculating the contig coverage of gene bodies and also calculated the fraction of assembled exons based on current annotations. To calculate the contig coverage of each gene we used the RSeQC control package with the RefGene database for hg19. The RefGene database includes known human genes (both protein-coding and non-coding) from the NCBI RNA reference sequences collection (RefSeq). We calculated the fraction of exons assembled by using bedmap from the bedops toolkit. For more details on the steps of the process refer to the supplemental document.

We compared these two measures of transcriptome quality on assemblies created without in silico normalization, Aspects I and II, and with in silico normalization Aspects I-Norm and II-Norm (Aspects 4 and 5 interchangeably). We saw a significant improvement in the coverage of RefGene gene bodies for assemblies created without in silico normalization (Figures 4a). For Aspect III, there was little difference in the coverage of RefGene gene bodies in any of the assemblies (Figure 4b). We observed a slight improvement of the fraction of Gencode exons assembled when comparing in silico normalization for Aspect I vs. Aspect I-Norm (Figure 5a and Table III). However, we observed a significant improvement of Gencode exons assembled (except at a downsampling of reads at 25% for Sample A) when comparing Aspect II vs. Aspect II-Norm (Figure 5a and Table III). For Aspect II vs. Aspect II-Norm Sample B, we observed a decreasing trend in p-value as the number of downsampled reads increased (Table III). For Aspect III, the largest differences in the assembly of Gencode exons was observed between the location of sequencing centers (Figure 5b). We compared the differences between assemblies performed from the SeQC dataset and a non-normalized assembly using >5.7 billion RNA-Seq reads from the Genotype-Tissue Expression (GTEx) Consortium. We found that the highest quality transcriptome assemblies were created from increasing the number of reads without normalization (Figure 6).

**Fig. 4:**
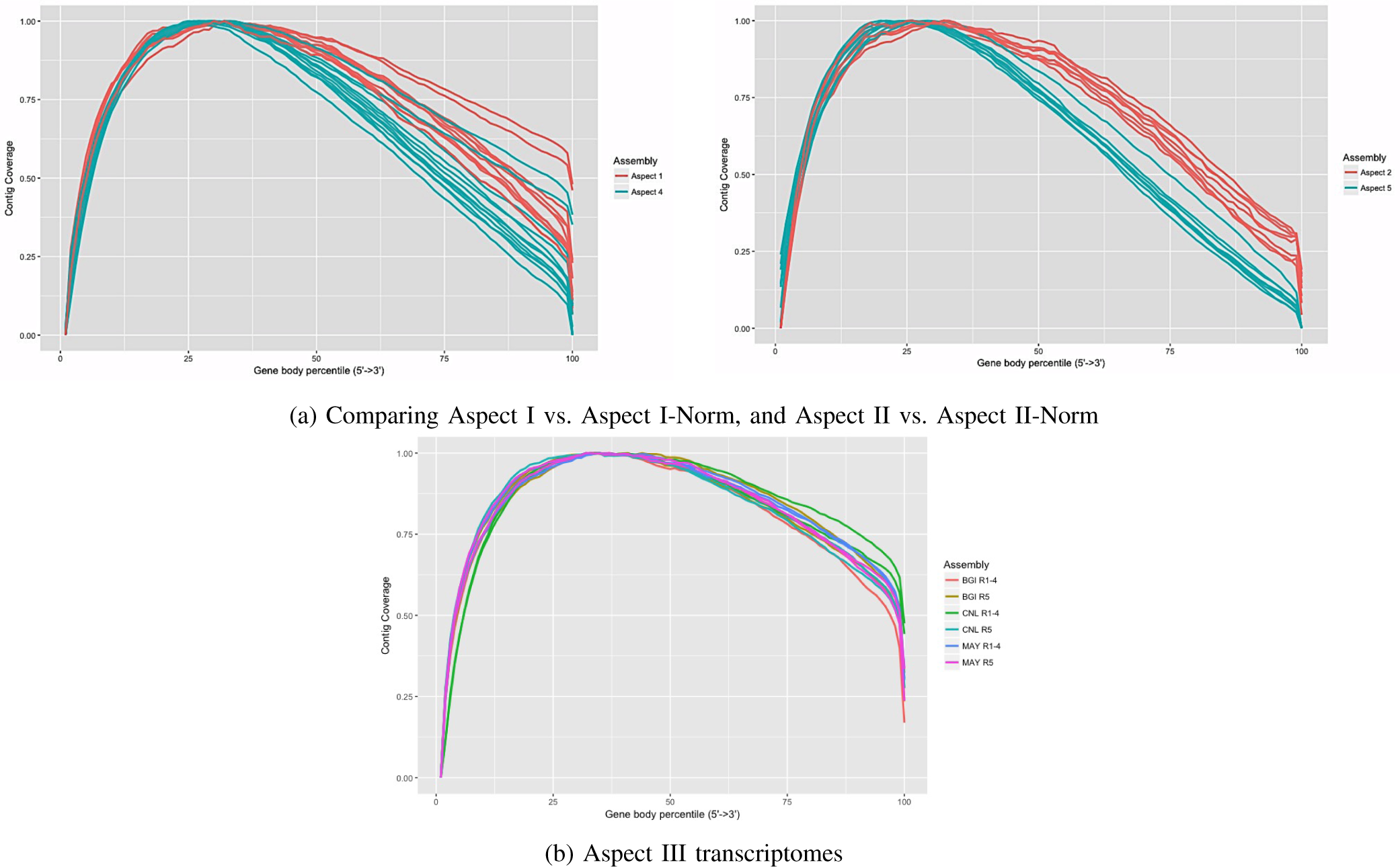
Contig coverage of gene body percentiles

**Fig. 5:**
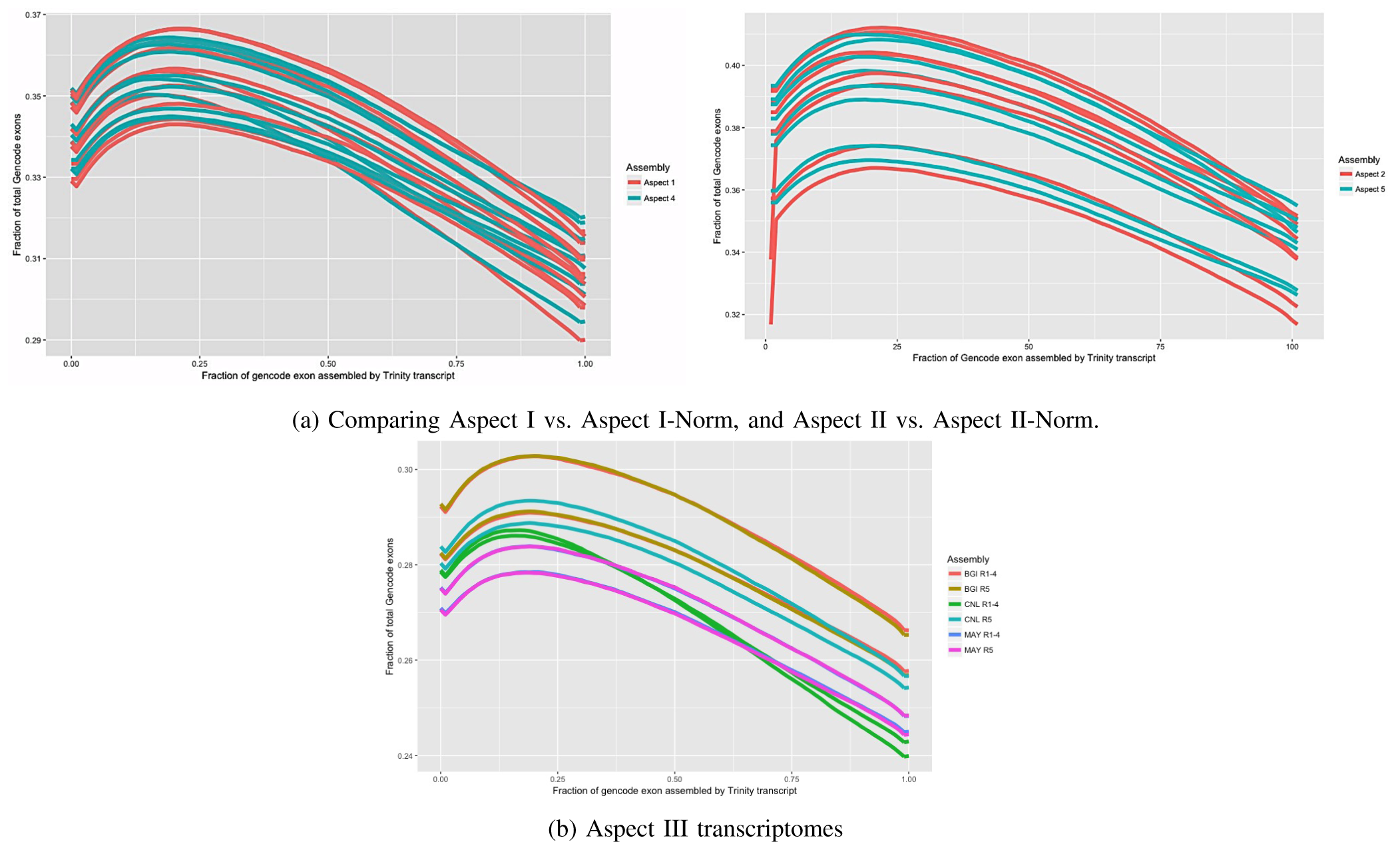
Fraction of Gencode exons assembled

**Fig. 6:**
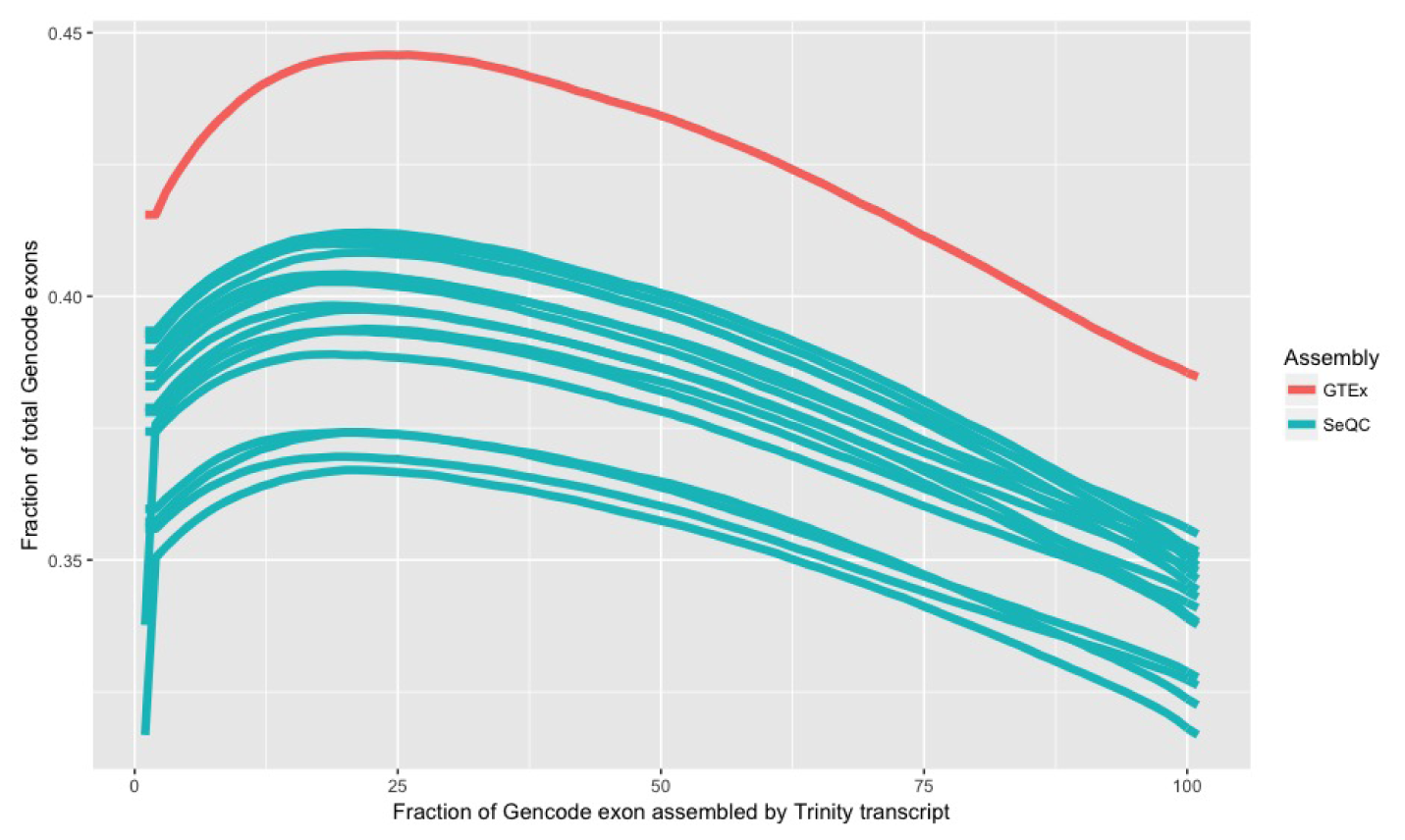
Fraction of Gencode exons assembled for comparing GTEx assembly vs. Assemblies from Aspect II and 5.

**Table III.**
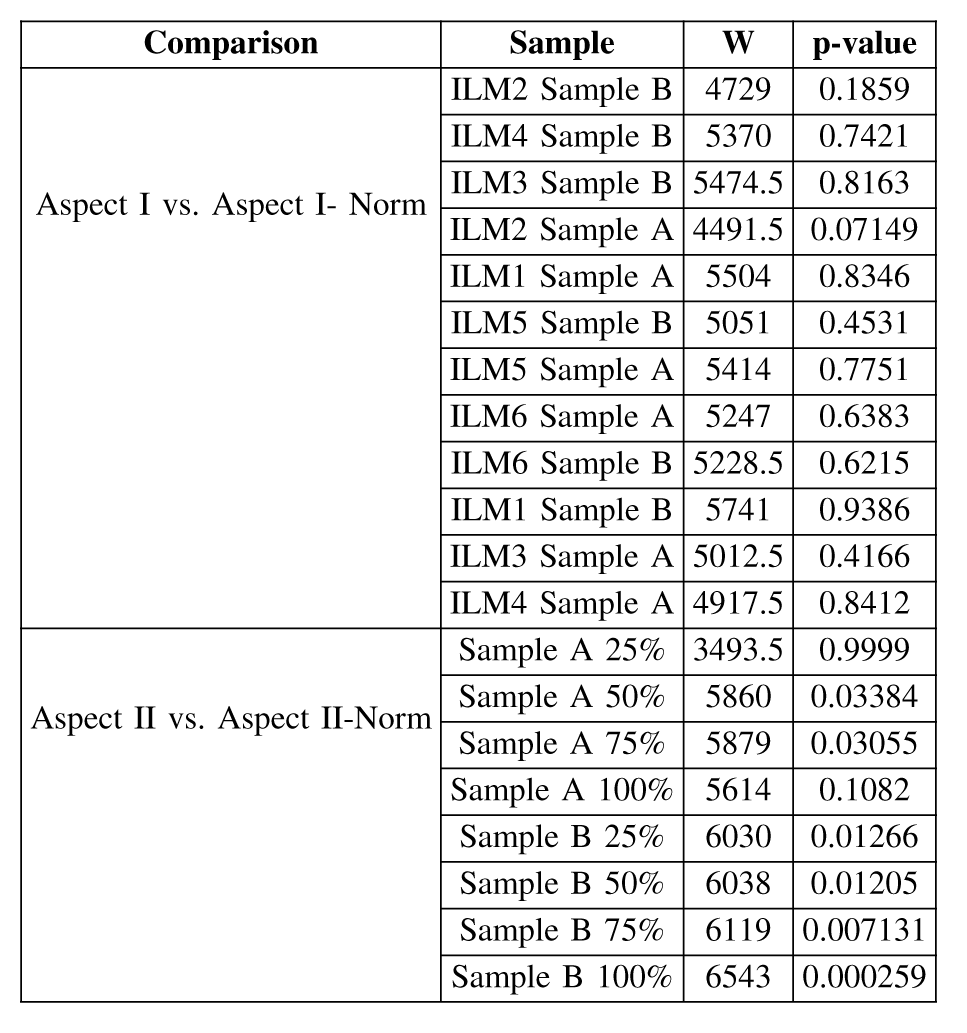
Mann-Whitney comparisons for all pairs of samples from Aspect I vs. Aspect I-Norm and Aspect II vs. Aspect II-Norm.

## IV. DISCUSSION

This study utilized reliable RNA-Seq data from the Sequencing Quality Control Consortium to examine the effects of library preparation, sequencing site and coverage on the transcriptome assembly. Our multi-step workflow consisted of pre-processing data to remove unnecessary attachments to the reads and to correct for the possible sequencing errors. The data was categorized into three aspects and assembled by Trinity tool. The resulting assembled transcriptomes went through a comprehensive quality control procedure to ensure the quality is satisfactory and also for comparing the assemblies.

The structural design and high volume RNA-Seq from SEQC study was instrumental in looking into the influence of sample handling on the output assembly. We verified the importance of the library preparation by showing that the high quality input data will generate more dependable transcriptome. The replicates from the sites that generated more reliable data performed equally well in transcriptome assembly as the vendor-prepared replicate. Our results also suggest that an aberrant coverage might be counterproductive in terms of assembly contiguity, but can improve gene body fraction. Therefore, if the resources are not a concern for data analysis, one might use more input data to have higher gene coverage. In summary, high quality library-preparation and appropriate genome coverage were the most important factors affecting the transcriptome assembly.

To emphasize the computational demands of the project, one should note that we successfully finished 52 assemblies through Aspects I, II, II, I-Norm and II-Norm for samples A and B, based on human RNA-Seq data. As Tables in the supplemental document shows the assembly runs range from 3X to 160X coverage of the human reference genome. As mentioned earlier, we computed the input read coverage based on reference genome for more accuracy and clarity. The proposed workflow comprises of three major components and the preprocessing steps were done once for each input dataset. This study was completed on the Blacklight, Greenfield and Bridges system at Pittsburgh Supercomputing Center (PSC). The computational aspects of runs on Blacklight and Greenfield system are provided at [28].

The size of the assembly jobs in this study exceeded many of the previous studies. The supercomputers at PSC enabled us to execute such large jobs. As an example of our very large runs, the Sample A 100 and Sample B 75 were two of highly resource demanding jobs, which were some of largest jobs ever completed on PSC. Both jobs ran simultaneously on a single 12 TB node on Bridges with 288 cores (HPE Integrity Superdome X with 16 Intel Xeon E7-8880 v3 CPUs 18 cores/CPU). Each job used half of the node, with access to up to 6 TB of RAM and 144 cores. Although Trinity does not generally scale to many cores (and can sometimes get slower with more threads), 4-6 TB of RAM was needed to complete these jobs. Tables II in the supplemental material shows the computational resources for these two long jobs.

Our experiments highlight the importance of the input data quality prior to sequencing. The sample libraries, prepared by laboratory personnel, highly impacts the accuracy of the transcriptome reconstruction, while sequencing with the same sequencing platform. Nonetheless, based on the different QC measures, the 3-5X coverage of reference genome should be sufficient for generating high quality transcriptome. The in-silico selection of high quality input (reads) also favorably benefits the outcome. Finally, we verified that de novo transcriptome assembly is reproducible among different sequencing sites, with the device causing minimal variability, compared to the profound human-related effects.

## ACKNOWLEDGMENT

This work used the Extreme Science and Engineering Discovery Environment (XSEDE), which is supported by National Science Foundation grant number ACI-1053575. Partial funding provided by Texas A&M AgriLife Research. We would like to thank the Epigenomics Core Facility at Weill Cornell Medicine, as well as the Starr Cancer Consortium (I9-A9-071), the Irma T. Hirschl and Monique Weill-Caulier Charitable Trusts, Bert L and N Kuggie Vallee Foundation, the WorldQuant Foundation, The Pershing Square Sohn Cancer Research Alliance, NASA (NNX14AH50G, NNX17AB26G), the National Institutes of Health (R21AI129851).

## REFERENCES

[1] Sanger, F., Nicklen, S., Coulson, A.R.: DNA sequencing with chain-terminating inhibitors. Proceedings of the National Academy of Sciences 74(12), 5463–5467 (1977)

[2] Voelkerding, K.V., Dames, S.A., D., D.J.: Next-generation sequencing: from basic research to diagnostics. Clinical Chemistry 55(4), 641–58 (2009)

[3] Grabherr, M.G., Haas, B.J., Yassour, M., Levin, J.Z., Thompson, D.A., Amit, I., Adiconis, X., Fan, L., Raychowdhury, R., Zeng, Q., et al.: Trinity: reconstructing a full-length transcriptome without a genome from RNA-seq data. Nature biotechnology 29(7), 644 (2011)

[4] Li, R., Zhu, H., Ruan, J., Qian, W., Fang, X., Shi, Z., Li, Y., Li, S., Shan, G., Kristiansen, K., et al.: De novo assembly of human genomes with massively parallel short read sequencing. Genome research 20(2), 265–272 (2010)

[5] Butler, J., MacCallum, I., Kleber, M., Shlyakhter, I.A., Belmonte, M.K., Lander, E.S., Nusbaum, C., Jaffe, D.B.: ALLPATHS: de novo assembly of whole-genome shotgun microreads. Genome research 18(5), 810–820 (2008)

[6] Shi, L., Reid, L.H., Jones, W.D., Shippy, R., Warrington, J.A., Baker, S.C., Collins, P.J., De Longueville, F., Kawasaki, E.S., Lee, K.Y., et al.: The microarray quality control (MAQC) project shows inter-and intraplatform reproducibility of gene expression measurements. Nature biotechnology 24(9), 1151–1161 (2006)

[7] Shippy, R., Fulmer-Smentek, S., et al.: Using rna sample titrations to assess microarray platform performance and normalization techniques. Nature biotechnology 24, 1123–1131 (2006)

[8] Baker, S.C., Bauer, S.R., Beyer, R.P., Brenton, J.D., Bromley, B., Burrill, J., Causton, H., Conley, M.P., Elespuru, R., Fero, M., et al.: The external RNA controls consortium: a progress report. Nature methods 2(10), 731–734 (2005)

[9] Consortium, SEQC/MAQC q. Iii., et al.: A comprehensive assessment of RNA-seq accuracy, reproducibility and information content by the sequencing quality control consortium. Nature biotechnology 32(9), 903–914 (2014)

[10] Kim, D., Pertea, G., Trapnell, C., Pimentel, H., Kelley, R., Salzberg, S.L., et al.: Tophat2: accurate alignment of transcriptomes in the presence of insertions, deletions and gene fusions. Genome Biol 14(4), 36 (2013)

[11] Thierry-MiegEmail, J. Danielle Thierry-MiegEmail: Aceview: a comprehensive cdna-supported gene and transcripts annotation. Genome Biology 7(s12) (2006)

[12] Glaus, P., Honkela, A., Rattray, M.: Identifying differentially expressed transcripts from rna-seq data with biological variation. Bioinformatics 28(13), 1721–1728 (2012)

[13] Liao, Y., Smyth, G.K., shi, W.S.: The subread aligner: fast, accurate and scalable read mapping by seed-and-vote. Nucleic Acids Research 4(10), 108 (2013)

[14] Dobin, A., Davis, C.A., Schlesinger, F., Drenkow, J., Zaleski, C., Jha, S., Batut, P., Chaisson, M., Gingeras, T.R.: Star: ultrafast universal rna-seq aligner. Bioinformatics 29(1), 15–21 (2013)

[15] Martin, M.: Cutadapt removes adapter sequences from high-throughput sequencing reads. EMBnet. journal 17(1), 10 (2011)

[16] Dodt, M., Roehr, J.T., Ahmed, R., Dieterich, C.: FLEXBAR - flexible barcode and adapter processing for next-generation sequencing platforms. Biology 1(3), 895–905 (2012)

[17] Le, H.-S., Schulz, M.H., McCauley, B.M., Hinman, V.F., Bar-Joseph, Z.: Probabilistic error correction for RNA sequencing. Nucleic acids research, 215 (2013)

[18] Wu, T.D., Watanabe, C.K.: Gmap: a genomic mapping and alignment program for mrna and est sequences. Bioinformatics 21(9), 1859–1875 (2005)

[19] McKenna, A., Hanna, M., Banks, E., Sivachenko, A., Cibulskis, K., Kernytsky, A., Garimella, K., Altshuler, D., Gabriel, S., Daly, M., et al.: The Genome Analysis Toolkit: a MapReduce framework for analyzing next-generation DNA sequencing data. Genome research 20(9), 1297–1303 (2010)

[20] Parra, G., Bradnam, K., Korf, I.: Cegma: a pipeline to accurately annotate core genes in eukaryotic genomes. Bioinformatics 23(9) (2007)

[21] Li, B., Fillmore, N., Bai, Y., Collins, M., Thomon, J.A., Stewart, R., Dewey, C.N.: Evaluation of de novo transcriptome assemblies from rna-seq data. Genome Biology 15(553) (2014)

[22] Ghaffari, N., Arshad, O.A., Jeong, H., Thiltges, J., Criscitiello, M.F., Yoon, B.-J., Datta, A., Johnson, C.D.: Examining de novo transcrip-tome assemblies via a quality assessment pipeline. IEEE/ACM Transactions on Computational Biology and Bioinformatics 1(1) (2015)

[23] Simaõ, F.A., Waterhouse, R.M., Ioannidis, P., Kriventseva, E.V., Zdobnov, E.M.: BUSCO: assessing genome assembly and annotation completeness with single-copy orthologs. Bioinformatics 31(19), 3210–3212 (2015)

[24] Li, B., Fillmore, N., Bai, Y., Collins, M., Thomson, J.A., Stewart, R., Dewey, C.N.: Evaluation of de novo transcriptome assemblies from rna-seq data. Genome Biol 15(12), 553 (2014)

[25] Abante, J., Ghaffari, N., Johnson, C.D., Datta, A.: Himme: using genetic patterns as a proxy for genome assembly reliability assessment. BMC Genomics 18(694) (2017)

[26] FastQC A Quality Control Tool for High Throughput Sequence Data. http://www.bioinformatics.babraham.ac.uk/projects/fastqc/

[27] Li, S., abaj, P.P., Zumbo, P., Sykacek, P., shi, W., shi, L., Phan, J., Wu, P.-Y., Wang, M., Wang, C., Thierry-Mieg, D., Jean, T.-M., Kreil, C.E. David P and Mason: Detecting and correcting systematic variation in large-scale rna sequencing data. Nature biotechnology 32, 888–895 (2014)

[28] Ghaffari, N., Abante, J., Blood, P.D., Singh, R., Johnson, C.D.: Computational considerations in transcriptome assemblies and their evaluation, using high quality human rna-seq data. In: Proceedings of the XSEDE16 Conference on Diversity, Big Data, and Science at Scale. doi:10.1145/2949550.2949572

